# Benchmarking Clustering Strategies for High-Dimensional Spike Time-Windows Data from Multi-Electrode Arrays

**DOI:** 10.64898/2026.09.17.752158

**Authors:** Lorenzo Sacchi, Sara Sommariva, Maurits Unkel, Femke M.S. de Vrij, Cristina Campi

## Abstract

From a statistical point of view, clustering spike time-windows from multi-electrode arrays (MEAs) recordings is a challenging high-dimensional, unsupervised clustering task where stationarity, commonly assumed in standard time-series analysis, is often violated and for which a gold standard is currently unavailable.

Here we aim at providing practical guidance on how to cluster this type of data by systematically comparing 108 clustering pipelines differing along three main dimensions: (i) the data feature space; (ii) the metrics used to quantify distance between data points; and (iii) the clustering algorithm. For our benchmark we used both labeled synthetic data, mimicking four physiologically-inspired classes of spike time-windows, and a set of real MEAs recordings from a two-dimensional in-vitro neural culture.

The performance of the competing pipelines was evaluated in terms of balanced accuracy and computational time when analyzing the synthetic datasets, while the Silhouette score was used for the real dataset, where no ground-truth is available. Overall, our analysis shows that the best combination is formed by k-means with Euclidean distance applied after Principal Component Analysis (PCA) of the spike time-windows. Conversely, hierarchical clustering showed the highest computational burden, while Independent Component Analysis and kernel PCA provided less effective noise suppression.

## 1. Introduction

Multielectrode Arrays (MEAs) represent a pivotal technological advance in neurophysiology, providing a non-invasive means to record in-vitro extracellular electrical signals over time from neuronal populations. This approach overcomes the severe technical and scaling limitations of traditional intracellular recording techniques, such as patch clamp, which are typically limited to a small number of simultaneously recorded neurons (Obien et al., 2015). The continuous development of this technology, from early designs to the emerging generation of high-density MEAs (HD-MEAs), has pushed the number of recording electrodes per sample into the thousands, creating a powerful tool for investigating complex network dynamics (Muthmann et al., 2015). At the same time, the scale and complexity of these recordings create substantial challenges for their statistical analysis and interpretation.

A first fundamental step in the analysis of MEAs data is the identification and grouping of extracellular action potentials, or spikes, detected in the time-series recorded by different sensors. This task typically involves extracting short time-windows around the detected spikes and clustering the resulting waveforms into groups that ideally correspond to distinct underlying neuronal sources (Hilgen et al., 2017). While extensive literature exists on clustering of time-series data across many domains (Aghabozorgi et al., 2015; Holder et al., 2024; Javed et al., 2020), clustering spike time-windows presents distinct challenges. Spikes are sharp, brief, low-amplitude events embedded in substantial background noise, and time-windows from temporally overlapping spikes generated by distinct neurons can superimpose, producing composite shapes that are difficult to disentangle. Furthermore, the number of underlying source generators and hence the appropriate number of clusters is typically unknown and can vary over the course of a recording. These factors, combined with the increasing spatial and time resolution of modern high-density MEAs, make spike time-windows clustering a challenging high-dimensional clustering problem for which a gold-standard procedure is still lacking (Ardelean and Portase, 2025).

Motivated by these considerations, this paper aims at designing a statistically robust and computationally efficient clustering pipeline specifically tailored for large scale spike time-windows datasets from MEAs recordings. Towards this end we systematically compared more than one hundred clustering pipelines that differ based on: (i) the representation of the data used for clustering, including both the original spike time-series and lower-dimensional derived feature representations; (ii) the dissimilarity measure used for quantifying distances between data points; (iii) the clustering algorithm. Benchmark datasets with ground-truth labels are currently not available for this problem, therefore we first evaluate the competing pipelines on a realistic synthetic datasets mimicking four physiologically-inspired classes of spike time-windows. This simulation study offers a controlled framework for assessing the impact of methodological choices on clustering performance and for identifying combinations of data representation, dissimilarity measures, and clustering algorithms that are effective in this context. The clustering pipeline that showed the best performance on synthetic data were then applied on real MEAs recording of a 2D neural culture to illustrate their use and interpretation in real settings.

The remaining of the paper is organized as follows: in Section 2 we summarize the methods we employed while in Section 3 we present the results we get on both synthetic and real data. Our discussion and conclusions are offered in Section 4.

## 2. Methods

MEAs electrodes can be spatially arranged in several configurations. In our case they are arranged in a square grid of size *S × S*, being *S*^2^ the total number of sensors in the grid. A first, commonly employed dimensionality-reduction step consists in selecting only a subset of sensors of interest, e.g. only sensors whose recording display a certain number of spikes.

Hence, denoted with *F* the number of time samples and with *n*_*s*_ the number of analyzed sensors, from a mathematical point of view recordings from MEAs can be represented as a set of time-series

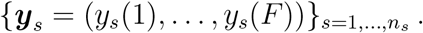

For each sensor *s*, we identify a set of *r*_*s*_ time-points, 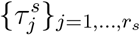, where spikes occur, resulting globally in a set of 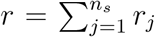 spike time-points. Spike detection is achieved by dividing ***y***_*s*_ in overlapping time-windows (length: 0.5 *s*, overlap: 0.25 *s*) and by identifying for each time-window the time-points where the signal exceeds *µ*_*w*_ *±* 5.4*σ*_*w*_, being *µ*_*w*_ and *σ*_*w*_ the mean and the median absolute deviation over the window, respectively.

After applying spike detection, for each spike *i ∈ {*1, *· · ·*, *r}*, denoted with *s*_*i*_ the sensor where such spike is detected, we extract from 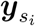 a time-window of length *p* centered at the spike-time point. In this way we build a dataset of *r p*-dimensional time-series, denoted as *{****x***_*i*_*}*_*i*=1,…,*r*_, where 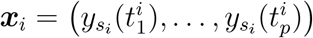, being 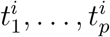 the time samples of the selected time-window.

Our objective is to define a robust and fast pipeline for partitioning the dataset *{****x***_*i*_*}*_*i*=1,…,*r*_ into *k* classes *C* = *{C*_1_, …, *C*_*k*_*}*. Clustering can be applied directly on the matrix ***X*** *∈* ℝ^*r×p*^ whose *i*-th row is the time-series ***x***_*i*_ or, alternatively, an additional step of dimensionality reduction can be applied projecting the dataset into a feature space of dimension *p*^*′*^ lower than *p*. Here, we test different combinations of dimensionality-reduction techniques, distance metrics, and clustering algorithms which are described in detail in the following sections.

### 2.1 Dimensionality reduction

Three possible dimensionality-reduction techniques are implemented: Principal Component Analysis (PCA, Pearson (1901)), a linear transformation seeking a set of orthogonal directions, the principal components, along which the variance of the data is maximized; Independent Component Analysis (ICA, Hyvarinen (1999)), a linear transformation seeking components that are mutually statistically independent; kernel PCA (Schölkopf et al., 1997) that maps the data into a high-dimensional reproducing kernel Hilbert space via a kernel function, and then performs standard PCA in that feature space. The result of these procedures is a new data matrix 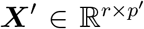 with *p*^*′*^ *< p* columns, that serves as input to the subsequent steps of our clustering pipeline. Before clustering, an initial evaluation of the effect of the dimensionality reduction can be assessed by backprojecting the reduced dataset into the original *p*-dimensional space and comparing it to the original time-series.

### 2.2 Distance metrics

Let **a** = (*a*_1_, …, *a*_*n*_) and **b** = (*b*_1_, …, *b*_*n*_) be two *n*-dimensional vectors representing two rows of the original data matrix ***X*** or the reduced one ***X***^*′*^, then the distance between **a** and **b** is quantified through the following metrics.

1. Minkowski distance (Minkowski, 1864) is defined as

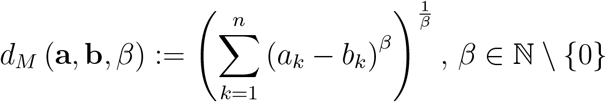

In our tests we considered *β* = 2 for which the Minkowski distance is the Euclidean distance (ED). The computational complexity of the Minkowski distance is *O*(*n*).
2. Pearson’s correlation coefficient distance (*ρ*_2_) (Golay et al., 1998) is defined as

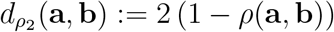

being *ρ*(**a, b**) the Pearson’s correlation coefficient between **a** and **b**. The computational complexity of the Pearson’s correlation coefficient distance is *O*(*n*).
3. Dynamic Time Warping (DTW) distance (Berndt and Clifford, 1994) and some of its variants. DTW distance, denoted as *d*_*DTW*_, is computed as the euclidean distance between **a** and **b** after realigning them in order to best match each other. This realignment relies on the pointwise distances *D*_*ij*_ = (*a*_*i*_ *− b*_*j*_)^2^ as shown in the Supplementary Material Web Algorithm 1. Derivative Dynamic Time Warping (DDTW) distance (Keogh and Pazzani, 2002) is obtained as the DTW distance between the first discrete derivatives of **a** and **b**, that is

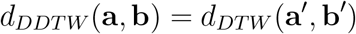

where 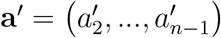 (and similarly **b**^*′*^) is a vector of length *n −* 2 whose entries are:

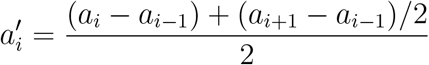

Weighted Dynamic Time Warping (WDTW) distance (Jeong et al., 2011) is computed as the DTW distance after applying a multiplicative weighting term to *D*_*ij*_ in order to penalize large shifts. Specifically we set, for all *i, j* = 1, …, *n*,

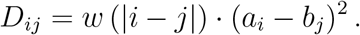

A classical choice for the weighting function is

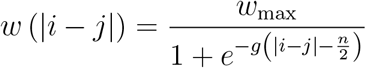

where *w*_*max*_ is a scale parameter representing the weights upper bound, while *g* defines the weights behavior, e.g. *g* = 0 represents constant weights, while *g* = 0.05 and *g* = 0.25 correspond to linear weights and sigmoidal weights, respectively (Jeong et al., 2011). In our analysis we set *w*_*max*_ = 2 and *g* = 0.25. Weighted Derivative Dynamic Time Warping (WDDTW) distance is defined as the WDTW distance computed between the first derivatives of **a** and **b**, namely

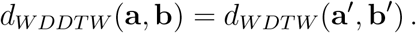

The computational complexity of the DTW distance (and variants) is *O*(*n*^2^).
4. Longest Common Sub-Sequence (LCSS) (Hirschberg, 1977) distance is defined as

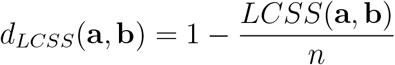

where *LCSS*(**a, b**) is the length of the longest (possibly non-contiguous) subsequence that appears in both vectors, computed as in Web Algorithm 2 of the Supplementary Material. The computational complexity of the LCSS distance is *O*(*n*^2^).
5. Edit distance on real sequences (EDR) (Chen et al., 2005) quantifies the dissimilarity between **a** and **b** as the minimum cost required to transform one into the other. The implemented algorithm for its computation is shown in the Supplementary Material, Web Algorithm 3. The computational complexity of the EDR distance is *O*(*n*^2^).
6. Short time series (STS) distance (Möller-Levet et al., 2003) quantifies the differences in the local changes of the two vectors and is defined as

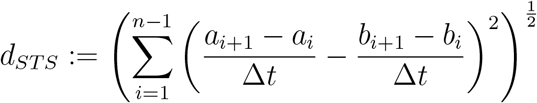

where Δ*t* is the distance between two consecutive time samples. The computational complexity of the STS distance is *O*(*n*).

### 2.3 Clustering algorithms

The clustering techniques we consider are k-means (KM) (MacQueen, 1967), fuzzy c-means (FCM) (Bezdek et al., 1984) and agglomerative hierarchical clustering (HC) with complete linkage method (Zepeda-Mendoza and Resendis-Antonio, 2013). We apply them to both the original and the reduced data, testing the different distance metrics implemented, as described above in Sections 2.1 and 2.2. Fixed the number of classes *k*, KM assigns each of the *r* observed time-series to one and only one class among *{C*_1_, *· · ·*, *C*_*k*_*}*, so as to minimize within-class dissimilarities. This is achieved by associating each time-series ***x***_*i*_ with a membership vector **u**_*i*_ *∈ {*0, 1*}*^*k*^, whose components correspond to the *k* classes. All components are equal to zero except the *j*-th one, which is equal to 1 indicating that ***x***_*i*_ is assigned to *C*_*j*_. The membership vectors are computed through an iterative scheme that in this study is initialized by a k-means++ approach (Arthur and Vassilvitskii, 2007).

FCM extends KM by allowing for fuzzy memberships: each observation may belong to multiple classes with different degrees of membership. Each time-series ***x***_*i*_ is thus associated to a membership vector **u**_*i*_ *∈* [0, 1]^*k*^ summing to 1. Similarly to KM, FCM is based on an iterative scheme here initialized by the output of KM. A parameter *m >* 1, known as fuzzy weighting exponent, regulates the fuzziness of the partition. In our tests we set *m* = 2 which is a common choice in the absence of a priori information (Bezdek et al., 1984).

Agglomerative HC is a sequential approach where each observation is initially assigned to a different cluster. Then, at each step, the two closest clusters are merged together until all the observations belong to the same cluster or some stopping criterion is met, e.g. a predefined number of clusters has been reached. HC requires computing between cluster distances. In our tests we used the complete linkage method which is less affected by the presence of outliers and noise on the data: given two clusters *C*_*i*_ and *C*_*j*_ their distance is defined as

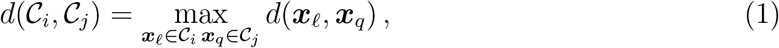

being *d* the chosen distance metric.

### 2.4 Selection of the number of clusters

For all the clustering algorithm, the number of clusters *k* is chosen using the Silhouette method (Rousseeuw, 1987). Given a partition *C* = *{C*_1_, …, *C*_*k*_*}*, for each time-series ***x***_*i*_, the Silhouette coefficient is defined as

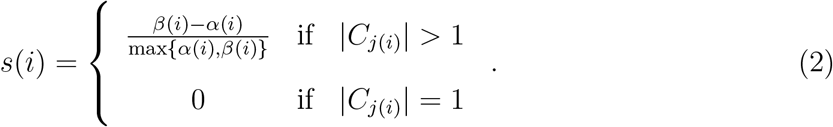

In Eq. (2), *C*_*j*(*i*)_ is the cluster to which ***x***_*i*_ belongs, |*C*_*j*(*i*)_| is its cardinality, while

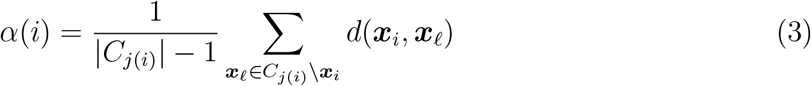

and

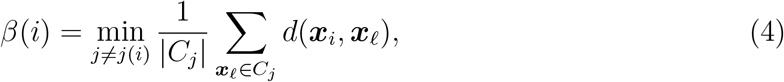

quantify, respectively, the distance of ***x***_*i*_ from its cluster and from all the other clusters, being *d* the distance metric employed in the clustering procedure. The Silhouette index, whose values range from *−*1 to 1, is computed as the average Silhouette coefficient across all the time-series in input:

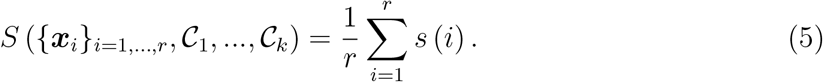

The optimal number of clusters is defined as the one corresponding to the highest Silhouette index observed within the evaluated cluster numbers.

## 3. Results

### 3.1 Synthetic dataset

In a first experiment we use synthetic spike time-series to compare the performance of the different clustering pipelines obtained by combining each one of the dissimilarity metrics with each one of the reduction and clustering techniques listed in the previous section. As shown in the first two rows of Figure 1, four physiologically-inspired spike classes are simulated: time-series in Class 1 are characterized by a narrow negative deflection followed by a small positive after-hyperpolarization; time-series in Class 2 feature a broad trough preceded by a small depolarizing pre-peak and succeeded by a prominent after-depolarization; time-series in Class 3 present a complex triphasic morphology, consisting of a positive onset, a sharp negative trough, and a positive rebound; time-series in Class 4 show a sharp positive leading peak followed by a shallow negative tail. For all classes, each time-series is constructed as a superposition of Gaussian basis functions defined over a time-window comprising *n* = 40 time samples, with the primary peak centered at the midpoint of the window. To introduce amplitude variability, each time-series is scaled by a random multiplicative factor uniformly drawn from [0.9, 1.1]. Additive Gaussian noise with zero mean and standard deviation equal to 0.05 is superimposed on each time-series to simulate background recording noise. A total of 100 spikes per class is simulated, yielding a balanced dataset of 400 time-series (Figure 1, third row).

**Figure 1.**
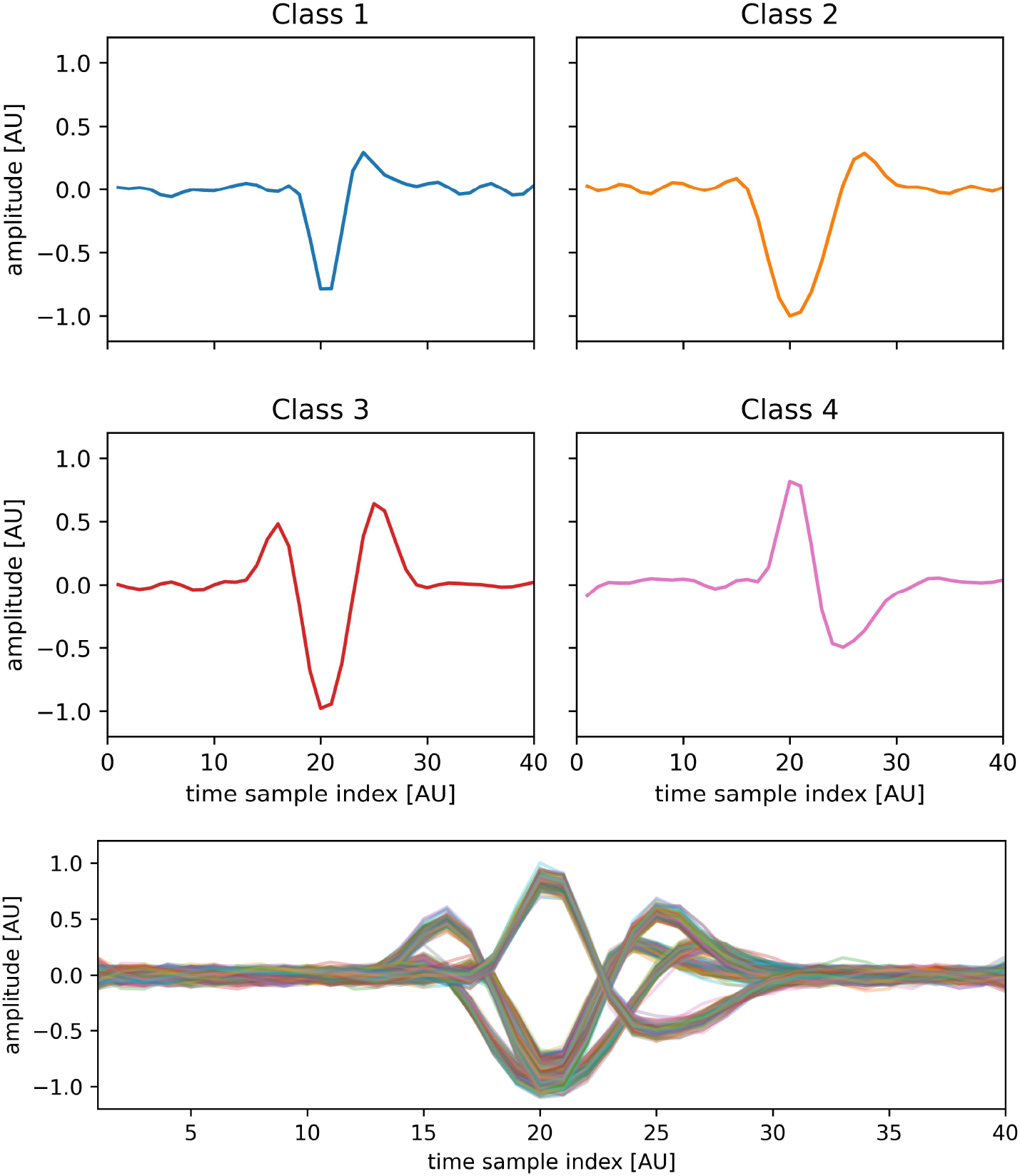
First two rows: illustrative time-series for the four physiologically-inspired simulated spike classes. Third row: whole synthetic dataset comprising 400 time-series.

The data are also reduced using PCA (retaining the components that explained at least 90% of the data variance), ICA (retaining 3 components) and kernel PCA (retaining 3 components). The number of components for ICA and kernel PCA is determined using the elbow method, plotting the mean squared error (MSE) between the original data and the backprojected one as a function of the number of retained components. The reduced datasets ***X***^*′*^ and their backprojections in the original *p*-dimensional space can be found in the Supplementary Material Web Figure 1.

We consider 6 as the maximum number of classes and we select the optimal number (*k*_*opt*_) using the Silhouette score as described in Section 2.4. As metric for the evaluation of the clustering results, we consider the balanced accuracy using the linear sum assignment algorithm (Kuhn, 1955) to find best alignment between classes and cluster labels.

The results of clustering are reported in the following as mean *±* standard deviation over 10 runs for KM and FCM, while just one run of HC is considered due to its deterministic nature. Specifically, Table 1 reports the estimated optimal number of clusters *k*_*opt*_, while the corresponding balanced accuracy are reported in Table 2. These tables show that, among the considered dimensionality-reduction techniques, the worst performance are obtained with kernel PCA which shows the highest variability, in terms of standard deviation, and the worst accuracy across runs. In the other cases, once the distance metric has been fixed, the three clustering algorithms perform similarly. Moreover, if clustering is performed on the original spike time-series dataset, all methods return the correct clusters (*k*_*opt*_ = 4 and balance accuracy equal to 1 for all runs) with all distance metrics, except DTW and WDTW. When the clustering algorithms are applied on the reduced dataset obtained with PCA or ICA, the correct clusters are obtained with ED, DTW, WDTW and STS.

**Table 1.** Number of clusters determined by the Silhouette method on noisy synthetic data.

| Optimal number of clusters ($k_{opt}$) | | | | | | | |
| --- | --- | --- | --- | --- | --- | --- | --- |
| KM |  | KM-PCA |  | KM-ICA |  | KM-kernelPCA |  |
| ED | $4.00 \pm 0.00$ | ED | $4.00 \pm 0.00$ | ED | $4.00 \pm 0.00$ | ED | $4.00 \pm 0.47$ |
| $\rho_2$ | $4.00 \pm 0.00$ | $\rho_2$ | $2.00 \pm 0.00$ | $\rho_2$ | $3.00 \pm 0.00$ | $\rho_2$ | $3.30 \pm 0.95$ |
| DTW | $3.00 \pm 0.00$ | DTW | $4.00 \pm 0.00$ | DTW | $4.00 \pm 0.00$ | DTW | $4.60 \pm 0.52$ |
| DDTW | $4.00 \pm 0.00$ | DDTW | $1.00 \pm 0.00$ | DDTW | $4.00 \pm 0.00$ | DDTW | $3.00 \pm 0.00$ |
| WDTW | $3.00 \pm 0.00$ | WDTW | $4.00 \pm 0.00$ | WDTW | $4.00 \pm 0.00$ | WDTW | $4.80 \pm 0.63$ |
| WDDTW | $4.00 \pm 0.00$ | WDDTW | $1.00 \pm 0.00$ | WDDTW | $4.00 \pm 0.00$ | WDDTW | $3.00 \pm 0.00$ |
| LCSS | $4.00 \pm 0.00$ | LCSS | $4.70 \pm 0.67$ | LCSS | $1.50 \pm 1.58$ | LCSS | $3.50 \pm 0.71$ |
| EDR | $4.00 \pm 0.00$ | EDR | $4.50 \pm 0.71$ | EDR | $1.00 \pm 0.00$ | EDR | $3.50 \pm 0.85$ |
| STS | $4.00 \pm 0.00$ | STS | $4.00 \pm 0.00$ | STS | $4.00 \pm 0.00$ | STS | $3.00 \pm 0.00$ |
| FCM |  | FCM-PCA |  | FCM-ICA |  | FCM-kernelPCA |  |
| ED | $4.00 \pm 0.00$ | ED | $4.00 \pm 0.00$ | ED | $4.00 \pm 0.00$ | ED | $3.90 \pm 0.32$ |
| $\rho_2$ | $4.00 \pm 0.00$ | $\rho_2$ | $2.00 \pm 0.00$ | $\rho_2$ | $3.00 \pm 0.00$ | $\rho_2$ | $3.10 \pm 0.32$ |
| DTW | $3.00 \pm 0.00$ | DTW | $4.00 \pm 0.00$ | DTW | $4.00 \pm 0.00$ | DTW | $4.80 \pm 0.42$ |
| DDTW | $4.00 \pm 0.00$ | DDTW | $1.00 \pm 0.00$ | DDTW | $4.00 \pm 0.00$ | DDTW | $3.00 \pm 0.00$ |
| WDTW | $3.00 \pm 0.00$ | WDTW | $4.00 \pm 0.00$ | WDTW | $4.00 \pm 0.00$ | WDTW | $4.90 \pm 0.32$ |
| WDDTW | $4.00 \pm 0.00$ | WDDTW | $1.00 \pm 0.00$ | WDDTW | $4.00 \pm 0.00$ | WDDTW | $3.00 \pm 0.00$ |
| LCSS | $4.00 \pm 0.00$ | LCSS | $4.70 \pm 0.82$ | LCSS | $1.00 \pm 0.00$ | LCSS | $3.10 \pm 0.57$ |
| EDR | $4.00 \pm 0.00$ | EDR | $4.50 \pm 0.85$ | EDR | $2.00 \pm 2.11$ | EDR | $3.10 \pm 0.32$ |
| STS | $4.00 \pm 0.00$ | STS | $4.00 \pm 0.00$ | STS | $4.00 \pm 0.00$ | STS | $3.00 \pm 0.00$ |
| HC |  | HC-PCA |  | HC-ICA |  | HC-kernelPCA |  |
| ED | 4.00 | ED | 4.00 | ED | 4.00 | ED | 4.00 |
| $\rho_2$ | 4.00 | $\rho_2$ | 2.00 | $\rho_2$ | 6.00 | $\rho_2$ | 3.00 |
| DTW | 3.00 | DTW | 4.00 | DTW | 4.00 | DTW | 3.00 |
| DDTW | 4.00 | DDTW | 1.00 | DDTW | 3.00 | DDTW | 3.00 |
| WDTW | 3.00 | WDTW | 4.00 | WDTW | 4.00 | WDTW | 3.00 |
| WDDTW | 4.00 | WDDTW | 1.00 | WDDTW | 3.00 | WDDTW | 3.00 |
| LCSS | 4.00 | LCSS | 1.00 | LCSS | 5.00 | LCSS | 1.00 |
| EDR | 4.00 | EDR | 1.00 | EDR | 5.00 | EDR | 1.00 |
| STS | 4.00 | STS | 4.00 | STS | 4.00 | STS | 4.00 |

**Table 2.**
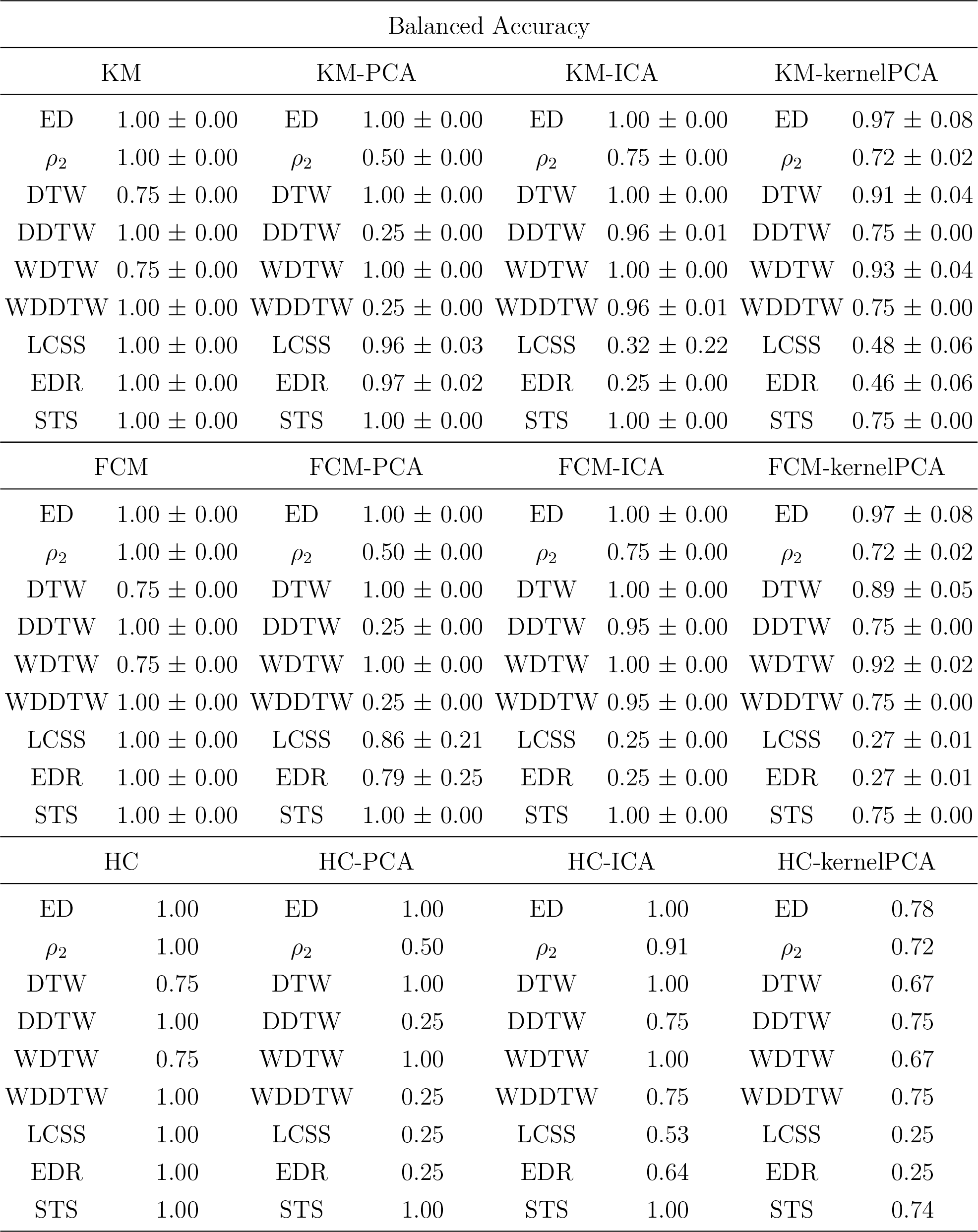
Clustering performance on noisy synthetic data with the optimal number of clusters determined by the Silhouette method.

| Balanced Accuracy |  |  |  |  |  |  |  |
| --- | --- | --- | --- | --- | --- | --- | --- |
| KM |  | KM-PCA |  | KM-ICA |  | KM-kernelPCA |  |
| ED | $1.00 \pm 0.00$ | ED | $1.00 \pm 0.00$ | ED | $1.00 \pm 0.00$ | ED | $0.97 \pm 0.08$ |
| $\rho_2$ | $1.00 \pm 0.00$ | $\rho_2$ | $0.50 \pm 0.00$ | $\rho_2$ | $0.75 \pm 0.00$ | $\rho_2$ | $0.72 \pm 0.02$ |
| DTW | $0.75 \pm 0.00$ | DTW | $1.00 \pm 0.00$ | DTW | $1.00 \pm 0.00$ | DTW | $0.91 \pm 0.04$ |
| DDTW | $1.00 \pm 0.00$ | DDTW | $0.25 \pm 0.00$ | DDTW | $0.96 \pm 0.01$ | DDTW | $0.75 \pm 0.00$ |
| WDTW | $0.75 \pm 0.00$ | WDTW | $1.00 \pm 0.00$ | WDTW | $1.00 \pm 0.00$ | WDTW | $0.93 \pm 0.04$ |
| WDDTW | $1.00 \pm 0.00$ | WDDTW | $0.25 \pm 0.00$ | WDDTW | $0.96 \pm 0.01$ | WDDTW | $0.75 \pm 0.00$ |
| LCSS | $1.00 \pm 0.00$ | LCSS | $0.96 \pm 0.03$ | LCSS | $0.32 \pm 0.22$ | LCSS | $0.48 \pm 0.06$ |
| EDR | $1.00 \pm 0.00$ | EDR | $0.97 \pm 0.02$ | EDR | $0.25 \pm 0.00$ | EDR | $0.46 \pm 0.06$ |
| STS | $1.00 \pm 0.00$ | STS | $1.00 \pm 0.00$ | STS | $1.00 \pm 0.00$ | STS | $0.75 \pm 0.00$ |
| FCM |  | FCM-PCA |  | FCM-ICA |  | FCM-kernelPCA |  |
| ED | $1.00 \pm 0.00$ | ED | $1.00 \pm 0.00$ | ED | $1.00 \pm 0.00$ | ED | $0.97 \pm 0.08$ |
| $\rho_2$ | $1.00 \pm 0.00$ | $\rho_2$ | $0.50 \pm 0.00$ | $\rho_2$ | $0.75 \pm 0.00$ | $\rho_2$ | $0.72 \pm 0.02$ |
| DTW | $0.75 \pm 0.00$ | DTW | $1.00 \pm 0.00$ | DTW | $1.00 \pm 0.00$ | DTW | $0.89 \pm 0.05$ |
| DDTW | $1.00 \pm 0.00$ | DDTW | $0.25 \pm 0.00$ | DDTW | $0.95 \pm 0.00$ | DDTW | $0.75 \pm 0.00$ |
| WDTW | $0.75 \pm 0.00$ | WDTW | $1.00 \pm 0.00$ | WDTW | $1.00 \pm 0.00$ | WDTW | $0.92 \pm 0.02$ |
| WDDTW | $1.00 \pm 0.00$ | WDDTW | $0.25 \pm 0.00$ | WDDTW | $0.95 \pm 0.00$ | WDDTW | $0.75 \pm 0.00$ |
| LCSS | $1.00 \pm 0.00$ | LCSS | $0.86 \pm 0.21$ | LCSS | $0.25 \pm 0.00$ | LCSS | $0.27 \pm 0.01$ |
| EDR | $1.00 \pm 0.00$ | EDR | $0.79 \pm 0.25$ | EDR | $0.25 \pm 0.00$ | EDR | $0.27 \pm 0.01$ |
| STS | $1.00 \pm 0.00$ | STS | $1.00 \pm 0.00$ | STS | $1.00 \pm 0.00$ | STS | $0.75 \pm 0.00$ |
| HC |  | HC-PCA |  | HC-ICA |  | HC-kernelPCA |  |
| ED | 1.00 | ED | 1.00 | ED | 1.00 | ED | 0.78 |
| $\rho_2$ | 1.00 | $\rho_2$ | 0.50 | $\rho_2$ | 0.91 | $\rho_2$ | 0.72 |
| DTW | 0.75 | DTW | 1.00 | DTW | 1.00 | DTW | 0.67 |
| DDTW | 1.00 | DDTW | 0.25 | DDTW | 0.75 | DDTW | 0.75 |
| WDTW | 0.75 | WDTW | 1.00 | WDTW | 1.00 | WDTW | 0.67 |
| WDDTW | 1.00 | WDDTW | 0.25 | WDDTW | 0.75 | WDDTW | 0.75 |
| LCSS | 1.00 | LCSS | 0.25 | LCSS | 0.53 | LCSS | 0.25 |
| EDR | 1.00 | EDR | 0.25 | EDR | 0.64 | EDR | 0.25 |
| STS | 1.00 | STS | 1.00 | STS | 1.00 | STS | 0.74 |

To assess the scalability of different analyses, we evaluate the computational cost of each combination of clustering algorithm, distance metric, and dimensionality-reduction approach as a function of the sample size. Specifically, we fix the number of clusters to 4 and we run each configuration with a sample size ranging from 10 (x1) to 400 (x40) time-series. At each sample size, time-series are drawn randomly from the original dataset so as to ensure approximately balanced classes. Results are depicted as the median elapsed time across 10 runs in Figure 2 for ED, *ρ*_2_, and DTW. Results for the other metrics are shown in the Supplementary Materials because STS behaves similar to ED, while DDTW, WDTW, WDDTW, LCSS and EDR are similar to DTW (Web Figures 3 and 4). For all metrics and clustering algorithms, the computational cost is strongly influenced by the choice of the dimensionality-reduction technique. Specifically, when ED and *ρ*_2_ are used (Figure 2, first and second rows respectively) kernel PCA shows the worst performance. Furthermore, for ED, when ICA and PCA are applied the elapsed time is similar to that obtained working in the original feature space, while for *ρ*_2_ PCA allows reducing the computational cost of almost two order of magnitude for all clustering algorithms. Instead, when DTW is used as distance metric (Figure 2, third row) all dimensionality-reduction techniques decrease the elapsed time with respect to the original feature space, PCA and ICA yielding the largest speedup (of two orders of magnitude). As far as clustering algorithms is concerned, in all cases HC shows the steepest growth while KM reach the best performances. Among the distance metrics, ED shows the overall lowest computational burden.

**Figure 2.**
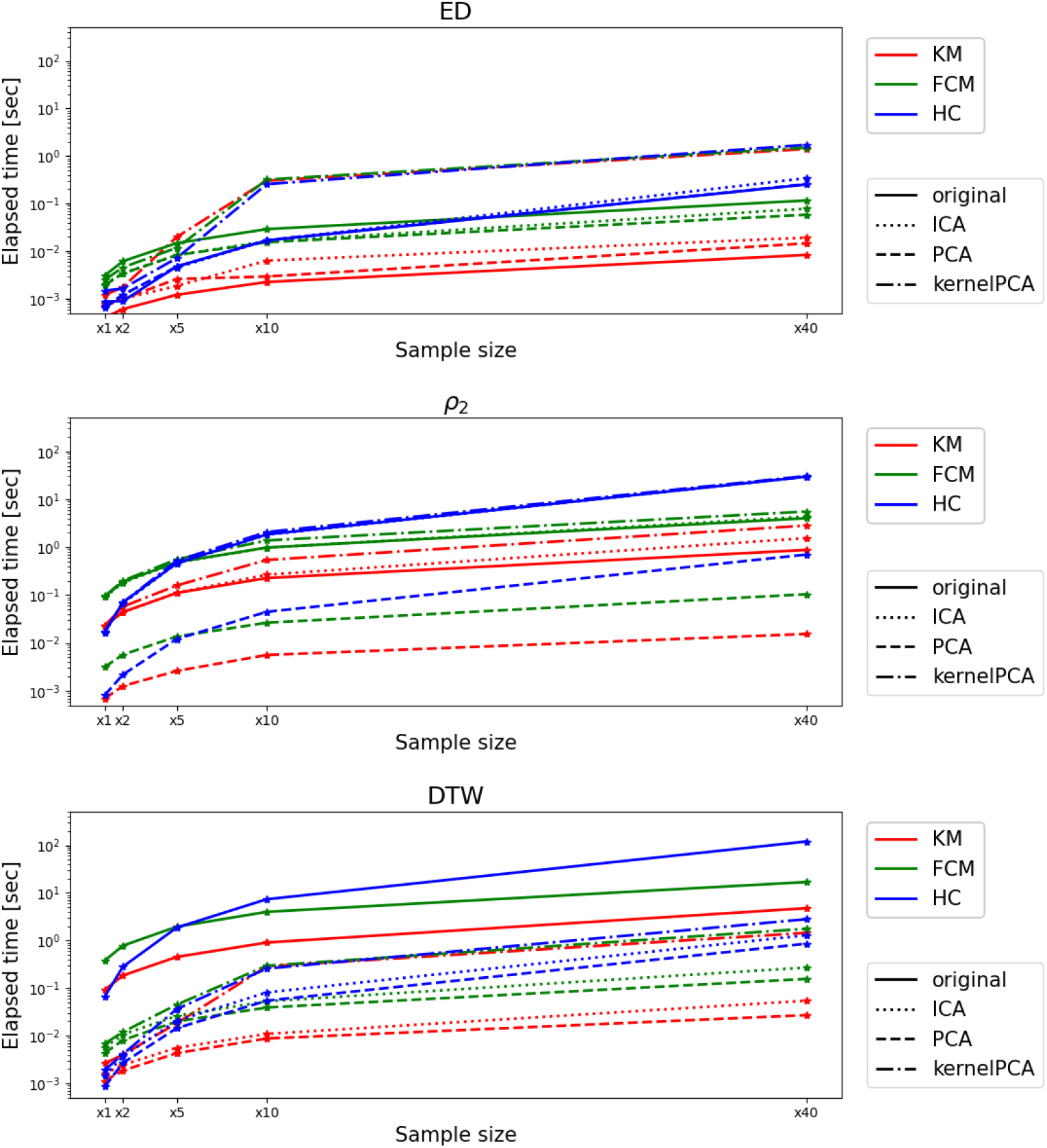
Computation cost for the different combinations of clustering approaches and dimensionality-reduction techniques when ED (first row), *ρ*^2^ (second row), and DTW (third row) are used as distance metrics. The median value across 10 different runs is displayed.

**Figure 3.**
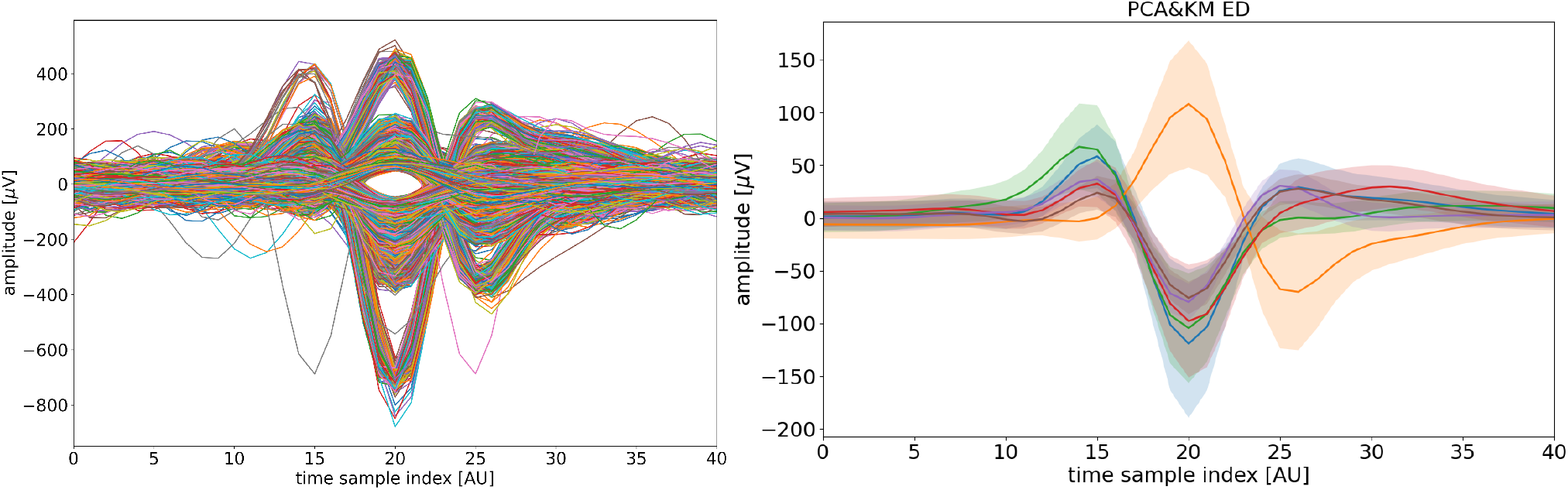
Best partition obtained from the analysis of the real data. Left: original spike time-series. Right: distribution of the clusters obtained applying PCA reduction techinque, K-means algorithm and Euclidean distance. For each cluster, the solid line depict its centroid, and the shaded area represents *±*1 standard deviation across cluster members at each time point.

Considering the above, we decided to apply to spike time-series from a real dataset only the following pipelines

- KM with ED and STS distance metrics applied to the original time-series;
- KM with ED, DTW, WDTW and STS distance metrics applied on the PCA reduced dataset;
- KM with ED, DTW, WDTW and STS distance metrics applied on the ICA reduced dataset.

### 3.2 Real dataset

The analysis uses a dataset built from a 3 minutes recording of a 2*D* human induced pluripotent stem cell (iPSC)-derived neural network culture that was acquired by seeding 3000 neural progenitor cells (NPCs) in MEA plates (3Brains CorePlate™) coated with laminin and maintained for 79 days in vitro. The sampling frequency was 19.753 kHz. The raw data are pre-processed applying Notch (frequency = 50 *Hz*) and band-pass (lowcut = 300 *Hz*, highcut = 3000 *Hz*) filters and the spikes are detected for all the channels, using the procedure described in Section 2. Channels are defined as active if they show a mean firing rate, computed as the ratio of the number of detected spikes and the recording time expressed in seconds, bigger than 0.25 spikes*/s*. Out of 4096 channels, only 305 are active and we collect spike time-windows coming from them for a gran-total of *r* = 132 772 time-windows. As feature reduction techniques we only perform PCA and ICA. We don’t perform kernel PCA as it showed the worst performance on synthetic data in terms of both balance accuracy (see Table 2) and computational time (see Figure 2). The original dataset is shown in Figure 3, left panel. The reduced datasets are shown in the Supplementary Material, Web Figure 2: for PCA, 4 components explain 90% of the variance, while for ICA the elbow method, with kneedle algorithm, selects 1 components.

As seen above in Section 3.1, when analyzing synthetic data, the best clustering technique configurations resulted to be the ones with KM, applied using ED and STS (with all the considered feature spaces) and with DTW and WDTW (after PCA and ICA). For each configuration, the number of clusters *k* is varied in the range *k ∈ {*4, 5, …, 10*}*, and selected through the Silhoutte index as shown in Section 2.4. Due to the O(*r*^2^) complexity of its computation and the large size *r* of the dataset, the Silhoutte index is computed on a randomly selected subdataset comprising 1000 waveforms.

Table 3 reports the best Silhouette score and corresponding optimal *k* for each configuration tested. In this scenario, clustering applied after PCA yields better results than clustering applied to the original time-series and after ICA. In particular, the best Silhoutte index is reached when KM with ED is applied after PCA with a number of cluster equal to 6. The centroids and relative variances of the resulting clusters are shown in Figure 3.

**Table 3.** Comparison of clustering configurations. The Silhouette score is the only metric directly comparable across configurations using different distance metrics.

| Algorithm | Distance | Best Silhouette | Optimal $k$ |
| --- | --- | --- | --- |
| KM | ED | 0.449 | 4 |
| KM | STS | 0.440 | 4 |
| KM-PCA | ED | **0.452** | 6 |
| KM-PCA | STS | 0.441 | 5 |
| KM-PCA | DTW | 0.426 | 4 |
| KM-PCA | WDTW | 0.425 | 4 |
| KM-ICA | ED | 0.304 | 5 |
| KM-ICA | STS | 0.393 | 4 |
| KM-ICA | DTW | 0.223 | 4 |
| KM-ICA | WDTW | 0.224 | 5 |

Since the Silhoutte index is known to be biased towards a smaller number of clusters (Teng et al., 2025), we test the effectiveness of the estimated number of clusters by considering the two following evaluation metrics (Arbelaitz et al., 2013):

- Davies-Bouldin index (Davies and Bouldin, 1979): non-negative, lower is better. Measures the average similarity between each cluster and its most similar cluster.
- Calinski-Harabasz index (Caliński and Harabasz, 1974): higher is better. Measures the ratio of between-cluster dispersion to within-cluster dispersion.

These two indices, in their `scikit-learn` implementation, do not accept a custom metric, while Silhouette score was computed using the same distance metric adopted for clustering

(Euclidean or STS respectively). Davies-Bouldin and Calinski-Harabasz can not be used to compare configurations using different distance metrics.

The best configuration is KM-PCA with ED with *k* = 6, selected based on the following criteria derived from the results shown in the Supplementary Material Web Tables 1 and 2:

- Silhouette score of 0.452.
- Sharp deterioration of the Silhouette score at *k* = 8 (0.341), suggesting that *k* = 7 represents a natural upper bound.
- The best value in the range *k ∈ {*4, …, 7*}* for Davies-Bouldin index is 0.862 for *k* = 6.
- Balanced cluster size distribution, with no degenerate clusters (smallest cluster: 10.1%, largest: 26.1%).

As shown in Table 3, when performing PCA the Silhouette index is higher than with ICA for all the considered distances. Coherently PCA results in a more effective noise suppression especially in the tails of the spike time-windows as can be seen from Web Figure 2 in the Supplementary Material.

## 4. Discussion

In this work we provide evidence-based recommendations for clustering large-scale spike time-windows datasets from MEA recordings, an emerging technology capable of measuring the activity over time of large neural assemblies with an outstanding temporal resolution. To achieve this we considered three different clustering algorithms (KM, FCM, and agglomerative HC) combined with nine dissimilarity measures (ED, *ρ*_2_, STS, as well as six elastic distances specifically designed for handling time-series) and applied either on the original spike time-windows or on a reduced feature space obtained with PCA, ICA or kernel PCA. The Silhouette score was used for choosing the optimal number of clusters. These methods have been chosen based on recent literature reviews and by preferring classical clustering pipelines over deep-learning techniques due to their higher balance between accuracy and overall runtime, especially due to the high-dimensional datasets typically produced in our context (Sadowska and Gajowniczek, 2026).

When analyzing a ground-truth labeled synthetic dataset, the best performance in terms of balanced accuracy and computational runtime was reached by KM applied to the original time-windows with ED and STS, or the PCA/ICA-based reduced feature space with ED, STS, DTW and WDTW. Conversely, FCM and HC showed higher computational runtimes, while, among the dimensionality reduction techniques, kernel PCA showed the highest variability across runs. The best performing combinations were further tested on a real dataset comprising 132 772 spike time-windows extracted from MEAs recordings of a 2D human induced pluripotent stem cell (iPSC)-derived neural culture. In this experiment the highest Silhouette score, equal to 0.452, was reached by applying KM with ED on the PCA-reduced dataset. When clustering is applied on the original time-windows the Silhouette score remained comparable values, while after ICA it dropped below 0.393 for all the distance metrics. The superiority of PCA over ICA in our application can be explained by the different objectives of the two decompositions: PCA maximises variance and preserves Euclidean geometry, which is consistent with the K-means objective function; ICA maximises statistical independence of components, which does not necessarily align with geometric cluster separability. This may motivate the overall higher noise-supression ability of PCA.

From a methodological point of view, one of the main limitations of the current work is that clustering is performed based solely on the shape of the spike waveforms without accounting for the temporal information on when the spikes occurred. Future efforts will be devoted to combining these two sources of information, for instance by jointly modeling waveform shape and inter-spike interval or firing rate dynamics, which may improve cluster separability for neurons with similar waveform morphology but distinct temporal firing patterns. A related direction involves interpreting clustering results in terms of interdependency, or connectivity, between sensors, possibly exploiting compositional data analysis (Campi et al., 2026). Second, as already stated our benchmark focused on classical clustering pipelines, excluding deep-learning-based approaches due to their higher computational cost and reduced interpretability in high-dimensional, large-scale settings. As increasingly efficient architectures become available, extending the comparison to include deep clustering methods represents a natural next step.

## Acknowledgements

This work was supported by the Horizon–EIC–2022–pathfinderopen project 3D–BrAIn (grant agreement 101098791). L.S., S.S., and C.C. are member of “Gruppo Nazionale per il Calcolo Scientifico” (INdAM-GNCS).

## Code and Data availability

The codes that implement the compared methods can be found at https://github.com/cristinacampi/3D-BrAIn_codes. The data underlying this article will be shared on reasonable request to the corresponding author.

## Notes

### Competing Interest Statement

The authors have declared no competing interest.

## References

Aghabozorgi, S., Shirkhorshidi, A. S., and Wah, T. Y. (2015). Time-series clustering–a decade review. Information systems 53, 16–38.

Arbelaitz, O., Gurrutxaga, I., Muguerza, J., Pérez, J. M., and Perona, I. (2013). An extensive comparative study of cluster validity indices. Pattern Recognition 46, 243–256.

Ardelean, E.-R. and Portase, R. L. (2025). A study of deep clustering in spike sorting. Neuroinformatics 23, 51.

Arthur, D. and Vassilvitskii, S. (2007). k-means++: the advantages of careful seeding. In Proceedings of the Eighteenth Annual ACM-SIAM Symposium on Discrete Algorithms, SODA ‘07, page 1027–1035, USA. Society for Industrial and Applied Mathematics.

Berndt, D. J. and Clifford, J. (1994). Using dynamic time warping to find patterns in time series. In Proceedings of the Third International Conference on Knowledge Discovery and Data Mining (AAAI WS’94), Seattle, WA, USA.

Bezdek, J. C., Ehrlich, R., and Full, W. (1984). FCM: The fuzzy c-means clustering algorithm. Computers and Geosciences 10, 191–203.

Caliński, T. and Harabasz, J. (1974). A dendrite method for cluster analysis. Communications in Statistics 3, 1–27.

Campi, C., Porro, F., Riccomagno, E., and Sommariva, S. (2026). A compositional framework for interpreting clustering of time series from neural cultures. In Conference of the International Federation of Classification Societies, pages 77–84. Springer.

Chen, L., Özsu, M. T., and Oria, V. (2005). Robust and fast similarity search for moving object trajectories. In Proceedings of the ACM SIGMOD International Conference on Management of Data, pages 491–502.

Davies, D. L. and Bouldin, D. W. (1979). A cluster separation measure. IEEE Trans. Pattern Anal. Mach. Intell. 1, 224–227.

Golay, X., Kollias, S., Stoll, G. D., Meier, A. V., and Boesiger, P. (1998). A new correlation-based fuzzy logic clustering algorithm for fMRI. Magnetic Resonance in Medicine 40, 249–260.

Hilgen, G., Sorbaro, M., Pirmoradian, S., Muthmann, J.-O., Kepiro, I. E., Ullo, S., Ramirez, C. J., Encinas, A. P., Maccione, A., Berdondini, L., et al. (2017). Unsupervised spike sorting for large-scale, high-density multielectrode arrays. Cell reports 18, 2521–2532.

Hirschberg, D. S. (1977). Algorithms for the longest common subsequence problem. Journal of the ACM 24, 664–675.

Holder, C., M. Middlehurst, and Bagnall, A. (2024). A review and evaluation of elastic distance functions for time series clustering. Knowledge and Information Systems 66, 765–809.

Hyvarinen, A. (1999). Fast and robust fixed-point algorithms for independent component analysis. Trans. Neur. Netw. 10, 626–634.

Javed, A., Lee, B. S., and Rizzo, D. M. (2020). A benchmark study on time series clustering. Machine Learning with Applications 1, 100001.

Jeong, Y.-S., Jeong, M. K., and Omitaomu, O. A. (2011). Weighted dynamic time warping for time series classification. Pattern Recognition 44, 2231–2240.

Keogh, E. J. and Pazzani, M. J. (2002). Derivative dynamic time warping. In Proceedings of the 1st SIAM International Conference on Data Mining (SDM), pages 239–241. SIAM.

Kuhn, H. W. (1955). The Hungarian method for the assignment problem. Naval Research Logistics Quarterly 2, 83–97.

MacQueen, J. B. (1967). Some methods for classification and analysis of multivariate observations. In Cam, L. M. L. and Neyman, J., editors, Proceedings of the Fifth Berkeley Symposium on Mathematical Statistics and Probability, volume 1 of Berkeley Symp. On Math. Statist. and Prob., pages 281–297, Berkeley, CA, USA. University of California Press.

Minkowski, H. (1864). Geometrie der Zahlen. Leipzig: Teubner.

Muthmann, J.-O., Amin, H., Sernagor, E., Maccione, A., Panas, D., Berdondini, L., Bhalla, U. S., and Hennig, M. H. (2015). Spike detection for large neural populations using high density multielectrode arrays. Frontiers in Neuroinformatics Volume 9 - 2015,.

Möller-Levet, C., Klawonn, F., Cho, K.-H., and Wolkenhauer, O. (2003). Fuzzy clustering of short time series and unevenly distributed sampling points. In Proceedings of the 5th International Symposium on Intelligent Data Analysis.

Obien, M. E. J., Deligkaris, K., Bullmann, T., Bakkum, D. J., and Frey, U. (2015). Revealing neuronal function through microelectrode array recordings. Frontiers in neuroscience 8, 423.

Pearson, K. (1901). Liii. on lines and planes of closest fit to systems of points in space. The London, Edinburgh, and Dublin Philosophical Magazine and Journal of Science 2, 559–572.

Rousseeuw, P. J. (1987). Silhouettes: A graphical aid to the interpretation and validation of cluster analysis. Computational and Applied Mathematics 20, 53–65.

Sadowska, M. and Gajowniczek, K. (2026). A comparative study of time series clustering performance with classification as a benchmark. Big Data and Cognitive Computing 10, 201.

Schölkopf, B., Smola, A., and Müller, K.-R. (1997). Kernel principal component analysis. In Gerstner, W., Germond, A., Hasler, M., and Nicoud, J.-D., editors, Artificial Neural Networks — ICANN’97, pages 583–588, Berlin, Heidelberg. Springer Berlin Heidelberg.

Teng, Z., Yan, J., Liu, D., and Zhang, P. (2025). When does the silhouette score work? a comprehensive study in network clustering.

Zepeda-Mendoza, M. L. and Resendis-Antonio, O. (2013). Use of metadata in visual interfaces to digital libraries. In Dubitzky, W., Wolkenhauer, O., Cho, K., and Yokota, H., editors, Encyclopedia of Systems Biology, pages 886–887. Springer, New York, NY.

