## Supplementary Material for "Benchmarking Clustering Strategies for High-Dimensional Spike Time-Windows Data from Multi-Electrode Arrays"

Lorenzo Sacchi[1], Sara Sommariva[1,*], Maurits Unkel[2], Femke M.S. de Vrij[2,3], Cristina Campi[1]
[1]Dipartimento di Matematica, Università di Genova, Italy
[2]Department of Psychiatry, Erasmus MC University Medical Center, Rotterdam, The Netherlands
[3]Erasmus MC Center of Expertise for Neurodevelopmental Disorders (ENCORE), Rotterdam, The Netherlands

#### Web Appendix A: Distance metrics implementation

This Web Appendix provides details on the implementation of some of the distances used in this work (DTW and its variant, Web Algorithm 1, LCSS distance, Web Algorithm 2, and EDR, Web Algorithm 3). Inspired by previous literature (?), for computing the LCSS and EDR distances we set $\varepsilon = \sigma/2$, being $\sigma$ the standard deviation of the data.

**Web Algorithm 1** DTW distance computation

**Initialize:** matrix $C$ of size $(n+1)\times(n+1)$ s.t. $C_{ij} = \begin{cases} 0 & \text{if } i=j=1 \\ +\infty & \text{otherwise} \end{cases}$
**for** $i = 1:n$ **do**
  **for** $j = 1:n$ **do**
    $D_{i,j} = (a_i - b_j)^2$
    $C_{i+1,j+1} = D_{i,j} + \min(C_{i,j}, C_{i,j+1}, C_{i+1,j})$
  **end for**
**end for**
**Return:** $d_{DTW} := C_{n+1,n+1}$

**Web Algorithm 2** LCSS distance computation

**Initialize:** matrix $L$ of size $(n+1)\times(n+1)$ s.t. $L_{i,j} = 0\ \forall i,j$.
**for** $i = 1:n$ **do**
  **for** $j = 1:n$ **do**
    **if** $|a_i - b_j| < \varepsilon$ **then**
      $L_{i+1,j+1} = L_{i,j} + 1$
    **else**
      $L_{i+1,j+1} = \max(L_{i,j+1}, L_{i+1,j})$
    **end if**
  **end for**
**end for**
$LCSS := L_{n+1,n+1}$
**Return:** $d_{LCSS} := 1 - LCSS/n$

**Web Algorithm 3** EDR distance computation

**Initialize:** matrix $E$ of size $(n+1) \times (n+1)$ s.t. $E_{i,j} = 0 \; \forall i, j$.
**for** $i = 1 : n$ **do**
  **for** $j = 1 : n$ **do**
    **if** $|a_i - b_j| < \varepsilon$ **then**
      $c = 0$
    **else**
      $c = 1$
    **end if**
    $match = E_{i,j} + c$
    $insert = E_{i,j+1} + 1$
    $delete = E_{i+1,j} + 1$
    $E_{i+1,j+1} = \min(match, insert, delete)$
  **end for**
**end for**
**Return:** $d_{EDR} := E_{n+1,n+1}$

### Web Appendix B: dimensionality reduction techniques evaluation

As described in the main text, a qualitative evaluation of the effect of the different dimensionality reduction techniques can be assessed by backprojecting the reduced dataset into the original $p$-dimensional space and comparing it to the original time-series. In this section we visualize such a comparison for the reduced datasets computed both from the synthetic (Web Figure 1) and the real (Web Figure ) dataset.

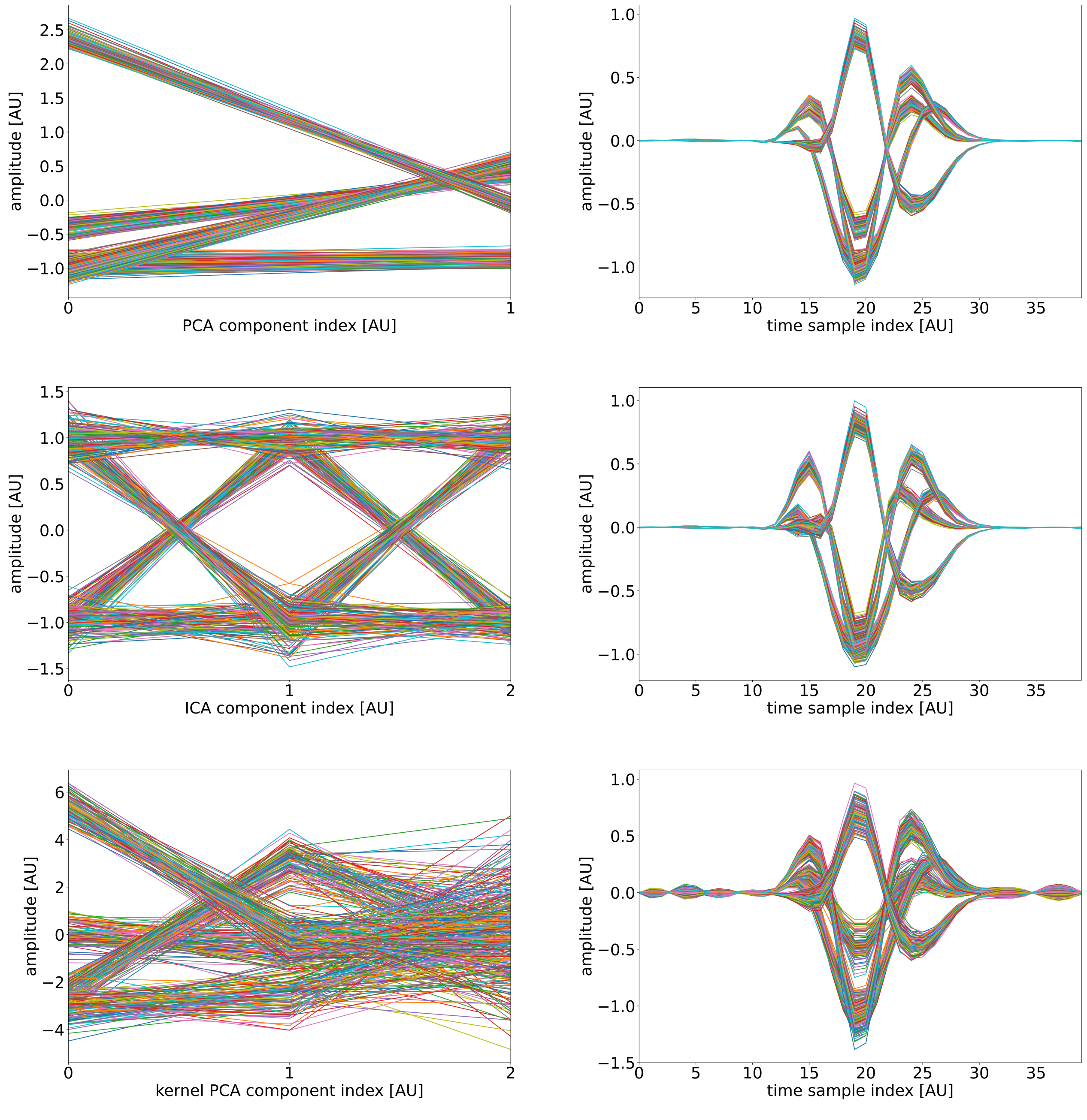


Web Figure 1: Synthetic dataset. First row: the data used for clustering post PCA (left panel), and the dataset transformed back using the 2 components (right panel). Second row: the data used for clustering post ICA (left panel), and the dataset transformed back using the 3 components (right panel). Third row: the data used for clustering post kernel PCA (left panel), and the dataset transformed back using the 3 components (right panel).

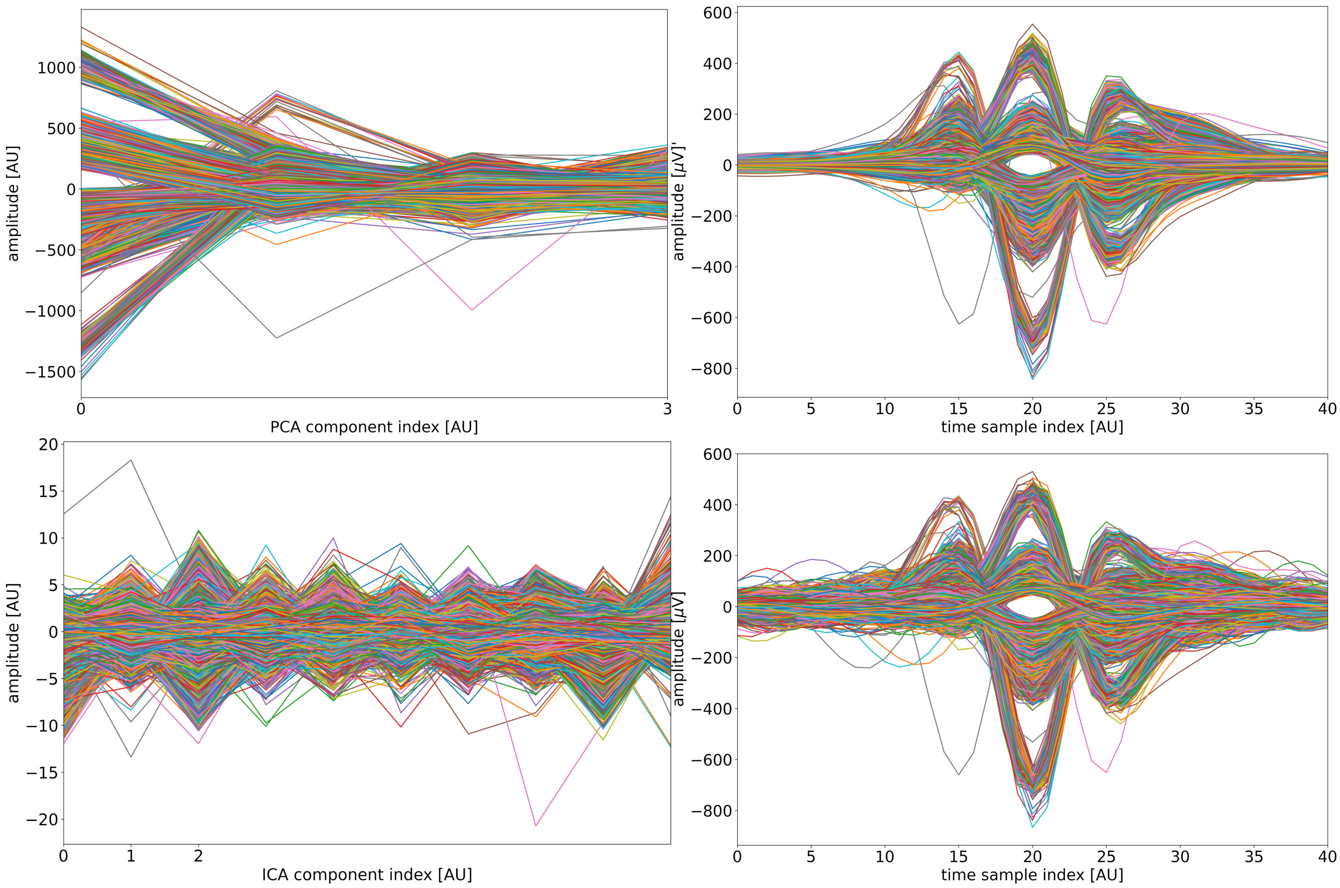


Web Figure 2: Real dataset. First row: the data used for clustering post PCA (left panel), and the dataset transformed back using the 2 components (right panel). Second row: the data used for clustering post ICA (left panel), and the dataset transformed back using the 3 components (right panel).

### Web Appendix C: Computational analysis for all metrics

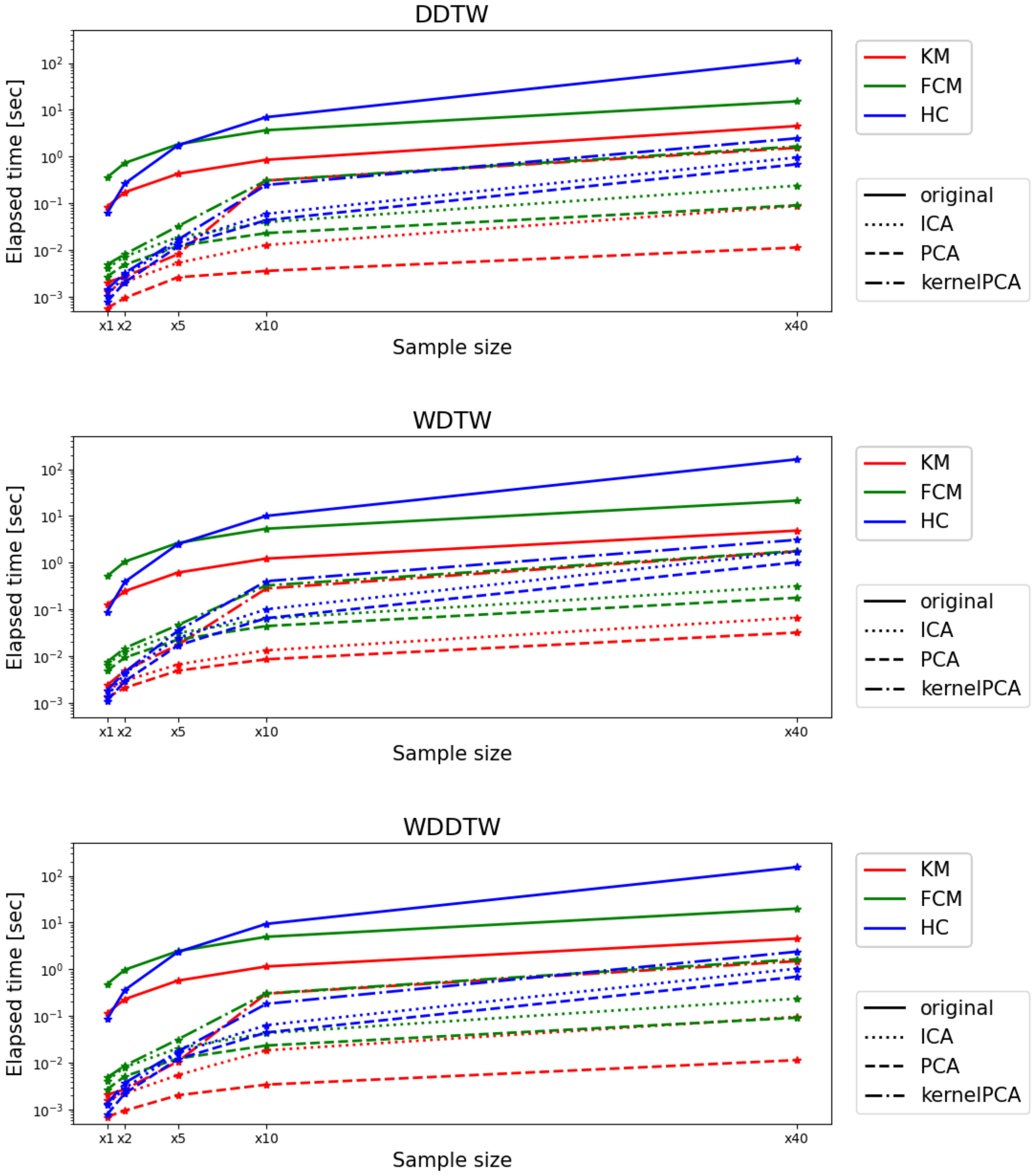


Web Figure 3: Computation cost for the different combinations of clustering techniques and feature spaces when dDTW (first row), wDTW (second row) and wdDTW (third row) and are used as distance metrics. The median value across 10 different runs is displayed.

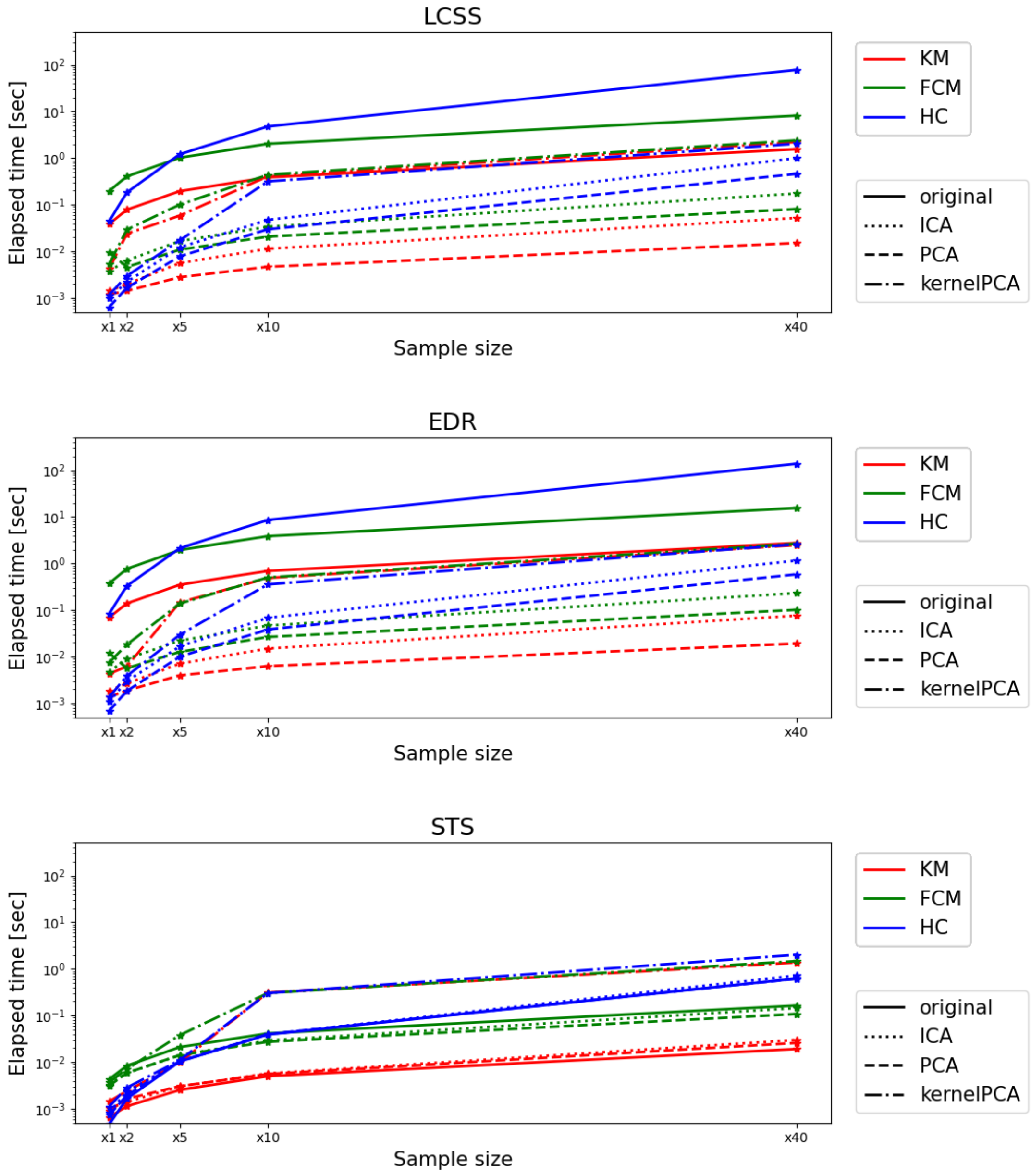


Web Figure 4: Computation cost for the different combinations of clustering techniques and feature spaces when LCSS (first row), EDR (second row) and STS (third row) and are used as distance metrics. The median value across 10 different runs is displayed.

### Web Appendix D: Additional tests on the real dataset.

Web Table 1: KM-PCA with ED: evaluation scores for $k \in \{4, \ldots, 10\}$.

| $k$ | Silhouette ↑ | Davies-Bouldin ↓ | Calinski-Harabasz ↑ |
|---|---|---|---|
| 4 | 0.449 | 0.930 | 202019 |
| 5 | 0.436 | 0.867 | 191123 |
| 6 | **0.452** | 0.862 | 182492 |
| 7 | 0.444 | 0.865 | 173251 |
| 8 | 0.341 | 0.950 | 163619 |
| 9 | 0.447 | 0.817 | 162463 |
| 10 | 0.445 | 0.846 | 156474 |

Web Table 2: Cluster size distribution for the optimal configuration ($k = 6$, KM-PCA, ED).

| **Cluster** | **Points** | **%** |
|---|---|---|
| 0 | 34589 | 26.1% |
| 1 | 34372 | 25.9% |
| 2 | 16144 | 12.2% |
| 3 | 13465 | 10.1% |
| 4 | 15917 | 12.0% |
| 5 | 18284 | 13.8% |

### References


Tan C.W., Herrmann M., S. M. e. a. (2025). Proximity forest 2.0: a new effective and scalable similarity-based classifier for time series. *Data Mining and Knowledge Discovery* **39,**.
